# A revised model for promoter competition based on multi-way chromatin interactions

**DOI:** 10.1101/612275

**Authors:** A. Marieke Oudelaar, Caroline L. Harrold, Lars L. P. Hanssen, Jelena M. Telenius, Douglas R. Higgs, Jim R. Hughes

## Abstract

Specific communication between gene promoters and enhancers is critical for accurate regulation of gene expression. However, it remains unclear how specific interactions between multiple regulatory elements and genes contained within a single chromatin domain are coordinated. Recent technological advances allow for the investigation of multi-way chromatin interactions at single alleles in individual nuclei. This can provide insights into how multiple regulatory elements cooperate or compete for transcriptional activation. We have used these techniques in a mouse model in which the α-globin domain is extended to include several additional genes. This allows us to determine how the interactions of the α-globin super-enhancer are distributed between multiple promoters in a single domain. Our data show that gene promoters do not form mutually exclusive interactions with the super-enhancer, but all interact simultaneously in a single complex. These finding show that promoters within the same domain do not structurally compete for interactions with enhancers, but form a regulatory hub structure, consistent with the recent model of transcriptional activation in phase-separated nuclear condensates.

## Introduction

An important question in current biology concerns the mechanisms by which genes are switched on and off during differentiation and development. Ultimately this is determined by interaction of the three fundamental regulatory elements of the genome: enhancers, promoters and boundary elements. The activity of these elements is closely related to the three-dimensional structure of the genome. Mammalian genomes are organized in topologically associating domains (TADs), which are self-interacting regions of chromatin, usually between 100 kb and 1 Mb in size (reviewed in Dekker and Heard, 2015; Dixon et al., 2016). The boundaries of TADs are often delineated by binding motifs for insulator proteins including CCCTC-binding factor (CTCF) and the promoters of actively transcribed genes (Rao et al., 2014; Sanyal et al., 2012). There is increasing evidence that TADs are formed by a process of active extrusion of chromatin loops which is limited by these boundary elements (Fudenberg et al., 2018; 2016; Sanborn et al., 2015).

The specificity of interactions between regulatory elements is dependent on the TAD structure of the genome. Enhancers preferentially interact with gene promoters in the same TAD (Symmons et al., 2014) and disruption of TAD boundaries results in promiscuous enhancer-promoter interactions and disrupted gene activity (de Wit et al., 2015; Guo et al., 2015; Hanssen et al., 2017; Lupiáñez et al., 2015; Narendra et al., 2015). However, it is not clear how specificity between multiple enhancer elements and promoters contained within a single TAD is regulated. Enhancers often exert different effects on what appear to be equally accessible genes within individual TADs. It has been proposed that enhancer-driven transcription from different promoters within a TAD is dependent on distance, orientation or affinity of the enhancers with respect to the specific promoters (Furlong and Levine, 2018; Long et al., 2016). Previous studies have suggested that enhancers may only interact with one accessible promoter at a time. This has led to a model in which the pattern of gene expression within a TAD containing multiple genes is determined by competition between promoters for limited access to shared enhancers. Based on this model, it has been proposed that co-expression of multiple genes regulated by shared enhancers in a single TAD results from rapidly alternating interactions of these genes with the enhancers in a “flip-flop” mechanism (Bartman et al., 2016; Wijgerde et al., 1995).

Recent evidence suggests that transcriptional activation takes place in nuclear condensates, which contain a high concentration of transcription factors, coactivators and components of the basal transcription machinery recruited by enhancer elements (Boehning et al., 2018; Boija et al., 2018; Cho et al., 2018b; Chong et al., 2018; Sabari et al., 2018). This implies that multiple enhancers within a TAD interact and function together in hub-like complexes, which have indeed been identified at a chromatin level (Allahyar et al., 2018; Oudelaar et al., 2018a). In the context of these recent findings, it is unclear if and how promoter competition occurs and what the underlying structural mechanism is.

We have recently developed Tri-C, a Chromosome Conformation Capture (3C) based approach which can analyze multi-way chromatin interactions at single alleles (Oudelaar et al., 2018a). Tri-C allows us to investigate whether promoters interact with enhancers in a mutually exclusive, one-to-one manner, or whether multiple promoters interact simultaneously with a shared set of enhancers in a hub-like structure. We have addressed this question using the well-characterized mouse α-globin locus as a model. The α-globin genes and their five enhancer elements, which fulfil the criteria for a super-enhancer (Hay et al., 2016), are located in a small TAD which is activated during erythroid differentiation (Brown et al., 2018). We have previously shown that *in vivo* deletion of two CTCF-binding sites at the upstream domain boundary results in an extension of the TAD and the incorporation of three upstream genes, which become highly upregulated under the influence of the strong α-globin super-enhancer (Hanssen et al., 2017). These mutant mice provide an excellent model to analyze the interactions between genes co-regulated by a set of well-characterized enhancers in primary cells.

By performing Tri-C in erythroid cells in which the CTCF boundary is deleted, we show that the upregulated gene promoters preferentially interact in hub-like complexes containing both the α-globin super-enhancer and the other active gene promoters in the domain. This shows that interactions between promoters and enhancers are not mutually exclusive and that there is no intrinsic structural competition between promoters for shared enhancers. These findings contribute to our understanding of the interplay between regulatory elements within and beyond TAD structures and the multiple layers or regulation that control gene expression.

## Results

We have previously defined all regulatory elements in and around the mouse α-globin cluster (Brown et al., 2018; Hanssen et al., 2017; Hay et al., 2016) (Fig. 1). The duplicated α-globin genes and the five globin enhancers (R1-R4 and Rm) lie within a small ~90 kb TAD. This TAD is flanked by predominantly convergent CTCF boundary elements. We have previously shown that deletion of the HS-38 and HS-39 CTCF-binding motifs causes strong upregulation of the upstream *Mpg*, *Rhbdf1* and *Snrnp25* genes in erythroid cells (Hanssen et al., 2017). To investigate how this deletion influences chromatin interactions with the α-globin super-enhancer, we performed Capture-C from the viewpoint of the strongest enhancer element, R2, in primary erythroid cells derived from wild type (WT) mice and mice in which the CTCF-binding sites were deleted (D3839). This showed an extension of the interaction domain in the D3839 mice, causing increased interactions between the α-globin enhancers and the *Mpg*, *Rhbdf1* and *Snrnp25* promoters (Fig. 1). The D3839 deletion thus creates an extended ~120 kb TAD in which the α-globin super-enhancer upregulates multiple genes. This extended domain enables us to address the mechanism by which a super-enhancer interacts with multiple accessible gene promoters in a single TAD.

**Figure 1:**
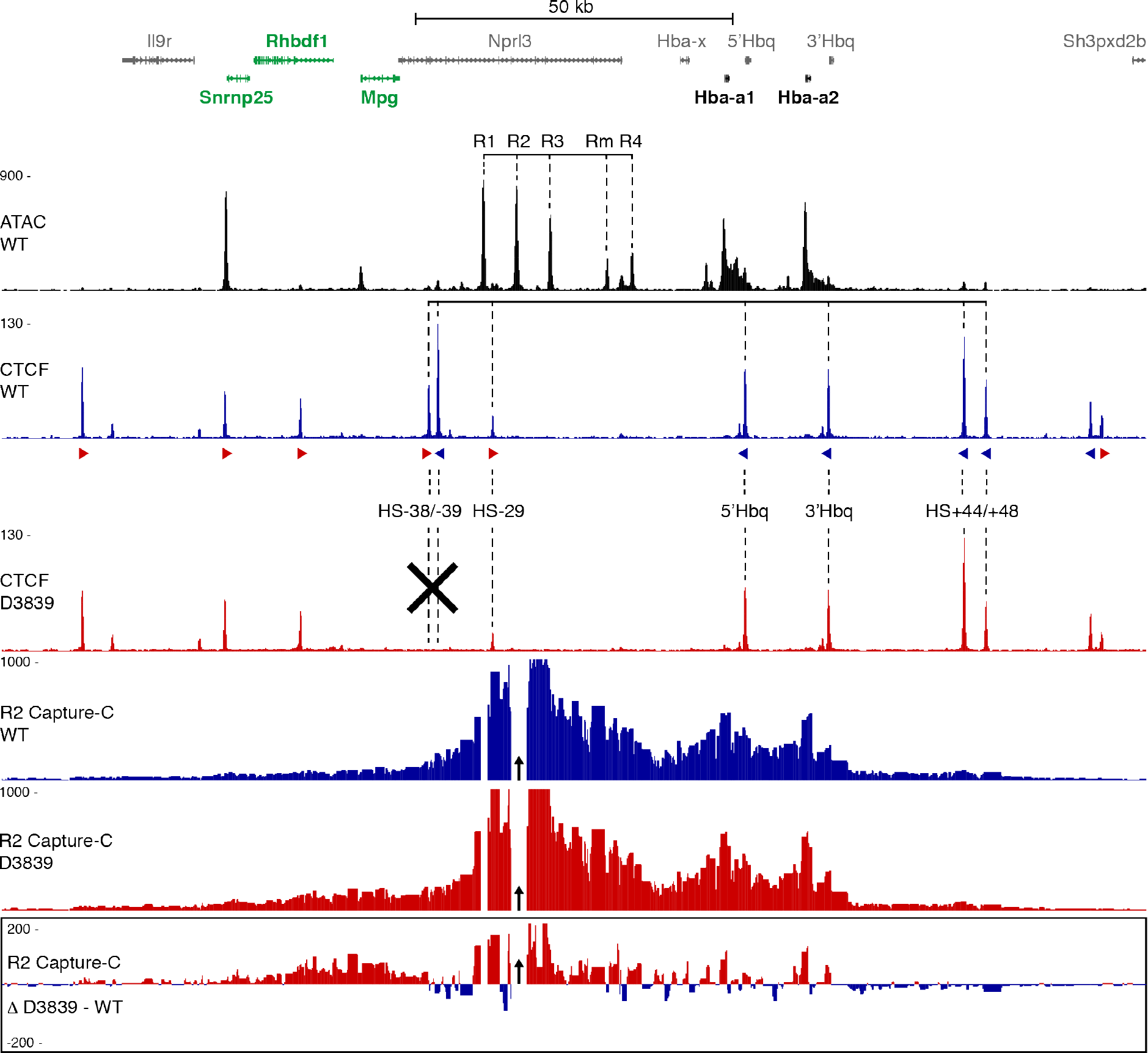
Characterization of a CTCF boundary deletion upstream of the α-globin locus. Gene annotation is shown at the top, with the α-globin genes in bold and genes upregulated by the deletion highlighted in green. Open chromatin (ATAC in WT erythroid cells) is shown below, with the α-globin enhancers highlighted. CTCF occupancy in WT (blue) and D3839 (red) erythroid cells is shown underneath, with the orientation of the CTCF binding motifs indicated by arrowheads (forward orientation in red; reverse orientation in blue). CTCF-binding sites of interested are highlighted and the deleted CTCF-binding sites are indicated with a black cross. The profiles below show Capture-C interactions from the viewpoint of the R2 enhancer (indicated with a black arrow) in WT (blue) and D3839 (red) erythroid cells, with a differential profile at the bottom. Profiles represent the mean number of normalized unique interaction counts per restriction fragment in 3 replicates. Coordinates (mm9): chr11:32,070,000–32,250,000.

Although Capture-C produces high-resolution Chromosome Conformation Capture (3C) profiles (Davies et al., 2015), it predominantly generates pair-wise interaction data. It is therefore not possible to determine the higher-order structures in which the multiple promoters and enhancers in the extended α-globin domain interact. Based on multi-way chromatin contacts generated by Tri-C, we have previously shown that the active α-globin locus is organized in a hub structure, in which multiple elements of the α-globin super-enhancer interact simultaneously with the α-globin promoters in a regulatory complex (Oudelaar et al., 2018a). To examine this structure in the context of the extended α-globin TAD containing multiple gene promoters, we performed a Tri-C experiment from the viewpoint of the R2 enhancer in primary D3839 and WT erythroid cells (Fig. 2 and Supplementary Fig. 2). Figure 2 shows the multi-way interactions detected by Tri-C. These are displayed in contact matrices in which we exclude the viewpoint of interest and plot the frequencies with which two elements interact simultaneously with this viewpoint in a single allele. Preferential, simultaneous interactions are visible as enrichments at the intersections between these elements, whereas mutually exclusive contacts between elements appear as depletions in the matrix. Consistent with our previous findings, we observe strong, simultaneous R2 interactions with the α-globin promoter and enhancer elements in WT cells. These interactions are not decreased in the D3839 cells, as would be expected if there was competition between these elements. Rather, there is a trend for increased interactions contributing to the α-globin hub in the D3839 cells, though this is not significant (Fig. 2b-c; green). In addition to the multi-way contacts between the α-globin enhancers and promoters, in D3839 we observe simultaneous interactions between the α-globin enhancer elements and the *Mpg* and *Rhbdf1* promoters (Fig. 2b-c, purple). Interestingly, the R2 contact matrix also shows simultaneous contacts with both the α-globin promoters and the *Mpg* and *Rhbdf1* promoters (Fig. 2b, grey). This indicates that all promoters in the extended D3839 TAD interact together in a single hub.

**Figure 2:**
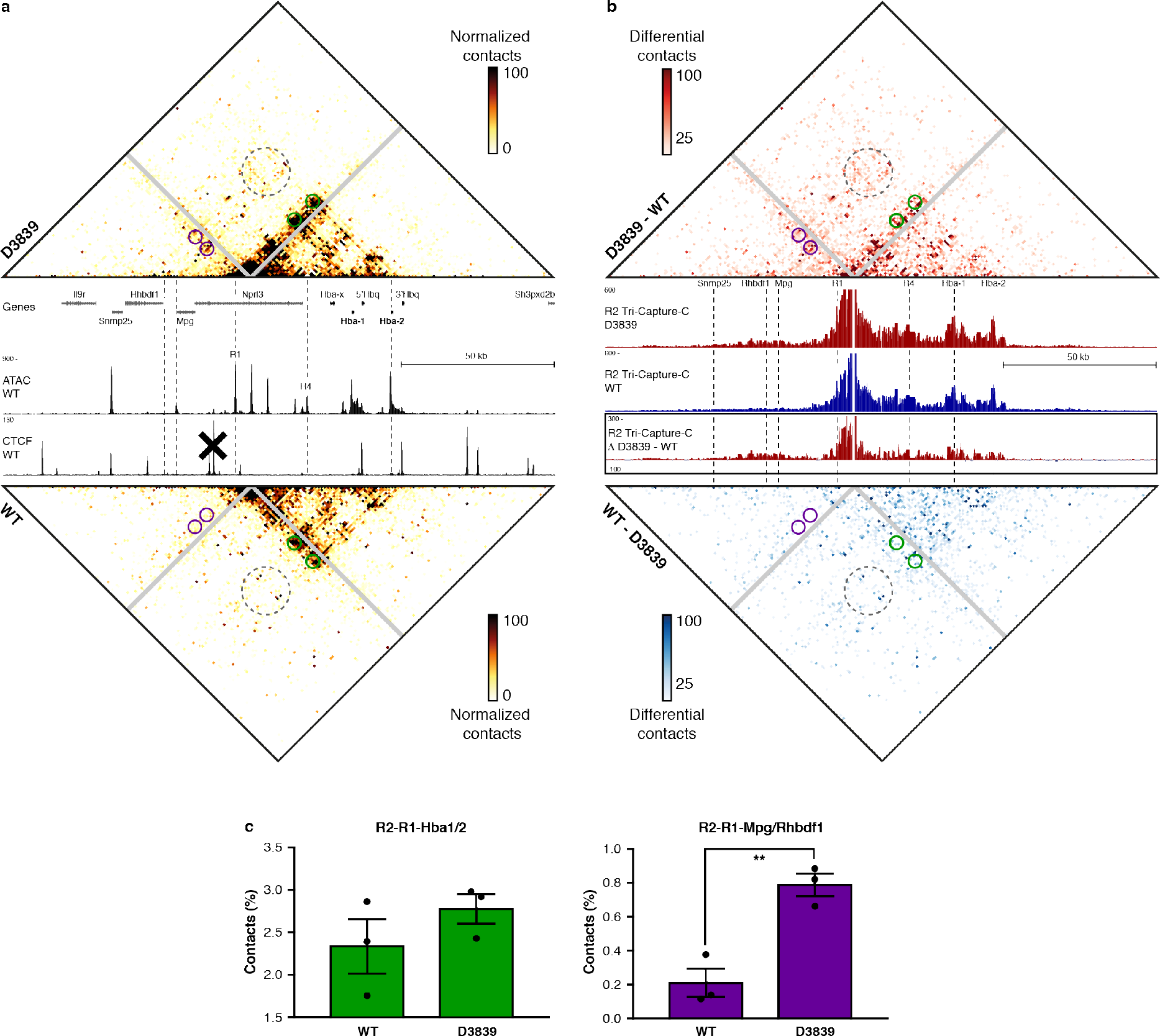
The formation of the enhancer-promoter hub at the α-globin locus is not dependent on the flanking CTCF boundary. **(a)** Tri-C contact matrices showing multi-way chromatin interactions with R2 in D3839 (top) and WT (bottom) erythroid cells. Matrices represent mean numbers of normalized, unique contact counts at 1 kb resolution in 3 replicates with proximity contacts around the R2 viewpoint excluded (gray diagonal). Gene annotation, open chromatin (ATAC) and CTCF occupancy in WT erythroid cells are shown in the middle. Coordinates (mm9): chr11:32,070,000–32,250,000. **(b)** Tri-C contact matrices showing differential multi-way chromatin interactions with R2 between D3839 and WT erythroid cells (top) and vice versa (bottom). Pair-wise interaction profiles derived from the Tri-C data from the R2 viewpoint (R2 Tri-Capture-C) are shown in the middle (D3839 in red, WT in blue), with a differential profile in the bottom panel. Coordinates (mm9): chr11:32,070,000–32,250,000. **(c)** Quantification of multi-way contacts between R2, R1, and the α-globin promoters (R2-R1-Hba1/2, green, P = 0.29)) and R2, R1, and the *Mpg* and *Rhbdf1* promoters (R2-R1-*Mpg*/*Rhbdf1*, purple, P = 0.0055). Quantified contacts are highlighted with corresponding colors in the matrices above. Numbers represent the proportion of these 3-way contacts relative to the total in the matrix and are averages of 3 replicates, with individual data points overlaid as dot plots and the standard error of the mean denoted by the error bar.

To allow more extensive examination of the simultaneous interactions occurring when the upstream genes interact with the α-globin super-enhancer in D3839 cells, we next generated Tri-C data from the viewpoint of the *Mpg* promoter (Fig. 3 and Supplementary Fig. 3). Comparison of multi-way *Mpg* interactions in D3839 and WT cells reveals a strong increase in interactions downstream of *Mpg* after removal of the CTCF boundary. These interactions are strongest proximal to the *Mpg* promoter and reduce in strength beyond the R1 enhancer, which is located close to the HS-29 CTCF-binding site. We also observe a clear increase in more distal downstream multi-way interactions, predominantly with the regions containing the α-globin promoters and enhancers. In a dynamic flip-flop model, the *Mpg* promoter would structurally compete with the α-globin promoters for dynamic interactions with the α-globin super-enhancer. Such mutually exclusive interactions would be reflected as a depletion of the corresponding multi-way interactions in the Tri-C matrix. However, we find preferential interactions between these elements. For example, when *Mpg* interacts with R1, it preferentially interacts with the α-globin promoters (Fig. 3, green) and R4 enhancer (Supplementary Fig. 3). Similarly, we find enrichment of multi-way interactions between *Mpg*, R1 and the *Rhbdf1* promoter (Fig. 3, purple). This shows that *Mpg* preferentially interacts with the α-globin super-enhancer in a complex that contains multiple enhancer elements and promoters. We also find that multi-way interactions between the three promoters are enriched (Fig. 3; orange), which further confirms that there is no structural competition between active promoters for contact with the super-enhancer within the extended TAD.

**Figure 3:**
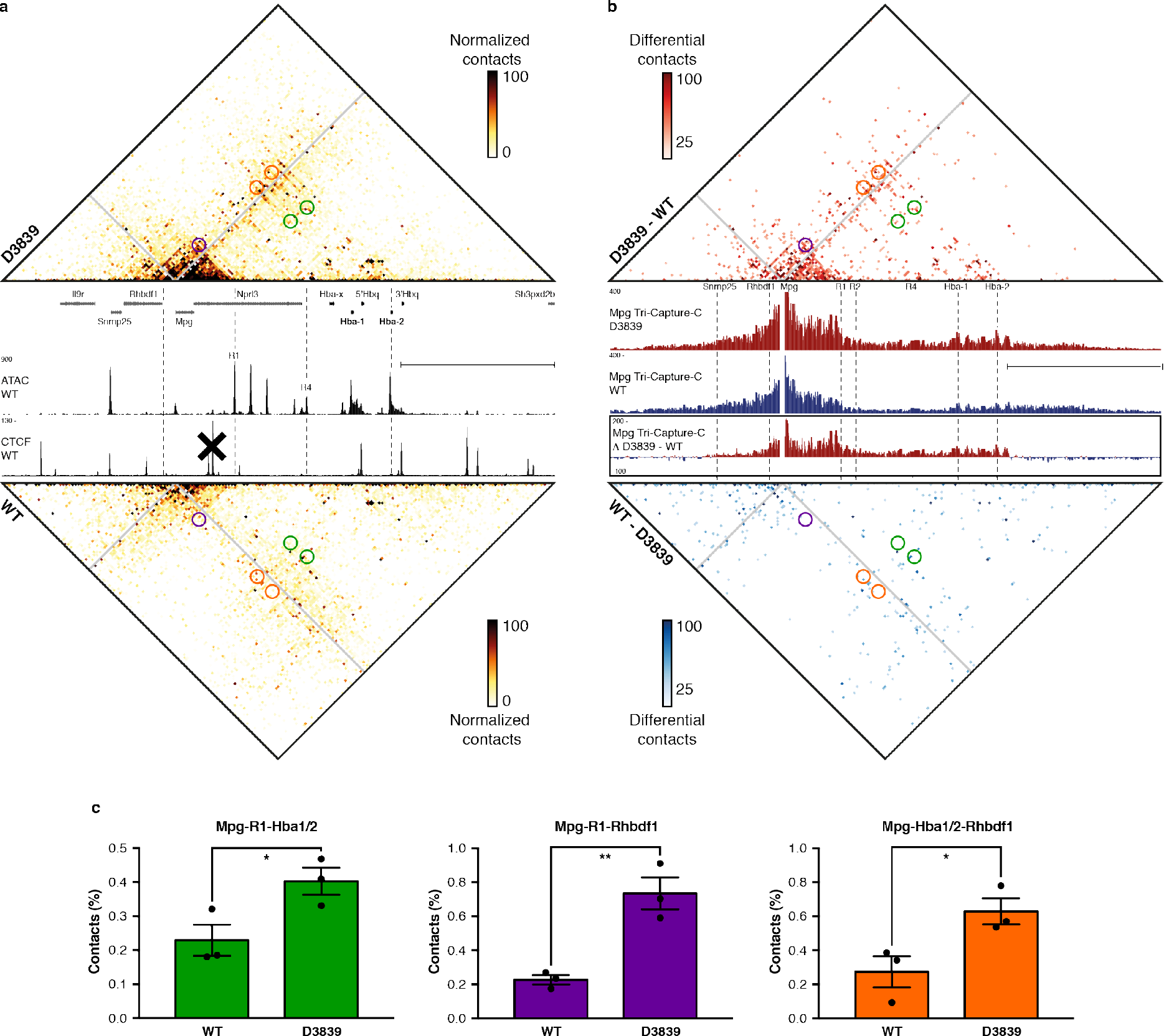
Deletion of a CTCF boundary results in the formation of a regulatory hub in which multiple gene promoters are incorporated. **(a)** Tri-C contact matrices showing multi-way chromatin interactions with *Mpg* in D3839 (top) and WT (bottom) erythroid cells. Matrices represent mean numbers of normalized, unique contact counts at 1 kb resolution in 3 replicates with proximity contacts around the *Mpg* viewpoint excluded (gray diagonal). Gene annotation, open chromatin (ATAC) and CTCF occupancy in WT erythroid cells are shown in the middle. Coordinates (mm9): chr11:32,070,000–32,250,000. **(b)** Tri-C contact matrices showing differential multi-way chromatin interactions with *Mpg* between D3839 and WT erythroid cells (top) and vice versa (bottom). Pair-wise interaction profiles derived from the Tri-C data from the *Mpg* viewpoint (*Mpg* Tri-Capture-C) are shown in the middle (D3839 in red, WT in blue), with a differential profile in the bottom panel. Coordinates (mm9): chr11:32,070,000–32,250,000. **(c)** Quantification of multi-way contacts between *Mpg*, R1 and the α-globin promoters (*Mpg*-R1-Hba1/2, green, P = 0.046); *Mpg*, R1 and the *Rhbdf1* promoter (*Mpg*-R1-*Rhbdf1*, purple, P = 0.0064); and *Mpg*, the α-globin promoters and the *Rhbdf1* promoter (*Mpg*-Hba1/2-*Rhbdf1*, orange, P = 0.040). Quantified contacts are highlighted with corresponding colors in the matrices above. Numbers represent the proportion of these 3-way contacts relative to the total in the matrix and are averages of 3 replicates, with individual data points overlaid as dot plots and the standard error of the mean denoted by the error bar.

## Discussion

To investigate how multiple regulatory elements and genes contained within a single TAD structurally interact, we analyzed multi-way chromatin interactions in an engineered extended TAD containing the α-globin super-enhancer and multiple gene promoters. We show that all gene promoters interact simultaneously with the five elements of the super-enhancer in a common regulatory hub. However, within the context of this extended TAD structure, the upstream non-globin genes do not interact as strongly and/or as frequently with the α-globin super-enhancer compared to the α-globin promoters. However, we show that when these genes form interactions with the α-globin enhancers, they preferentially interact in a complex in which the α-globin promoters are also present (Fig. 3, Supplementary Fig. 3). By comparing the α-globin hub in WT and D3839 cells, we show that the inclusion of additional promoters to this complex does not weaken the interactions between the α-globin promoters and enhancers and might even have an overall stabilizing effect on the hub (Fig. 2).

Our data thus show that multiple gene promoters can simultaneously interact with a shared super-enhancer at a single allele. This demonstrates that the previously reported “flip-flop” model of promoter competition, in which individual gene promoters interact with enhancers in a mutually exclusive manner, is not universally true and that there is no intrinsic competition between gene promoters for physical access to shared enhancers within a single TAD (Fig. 4). Our model is supported by recent live imaging experiments in Drosophila which showed coordinated bursting of two genes regulated by a single shared enhancer (Fukaya et al., 2016).

**Figure 4:**
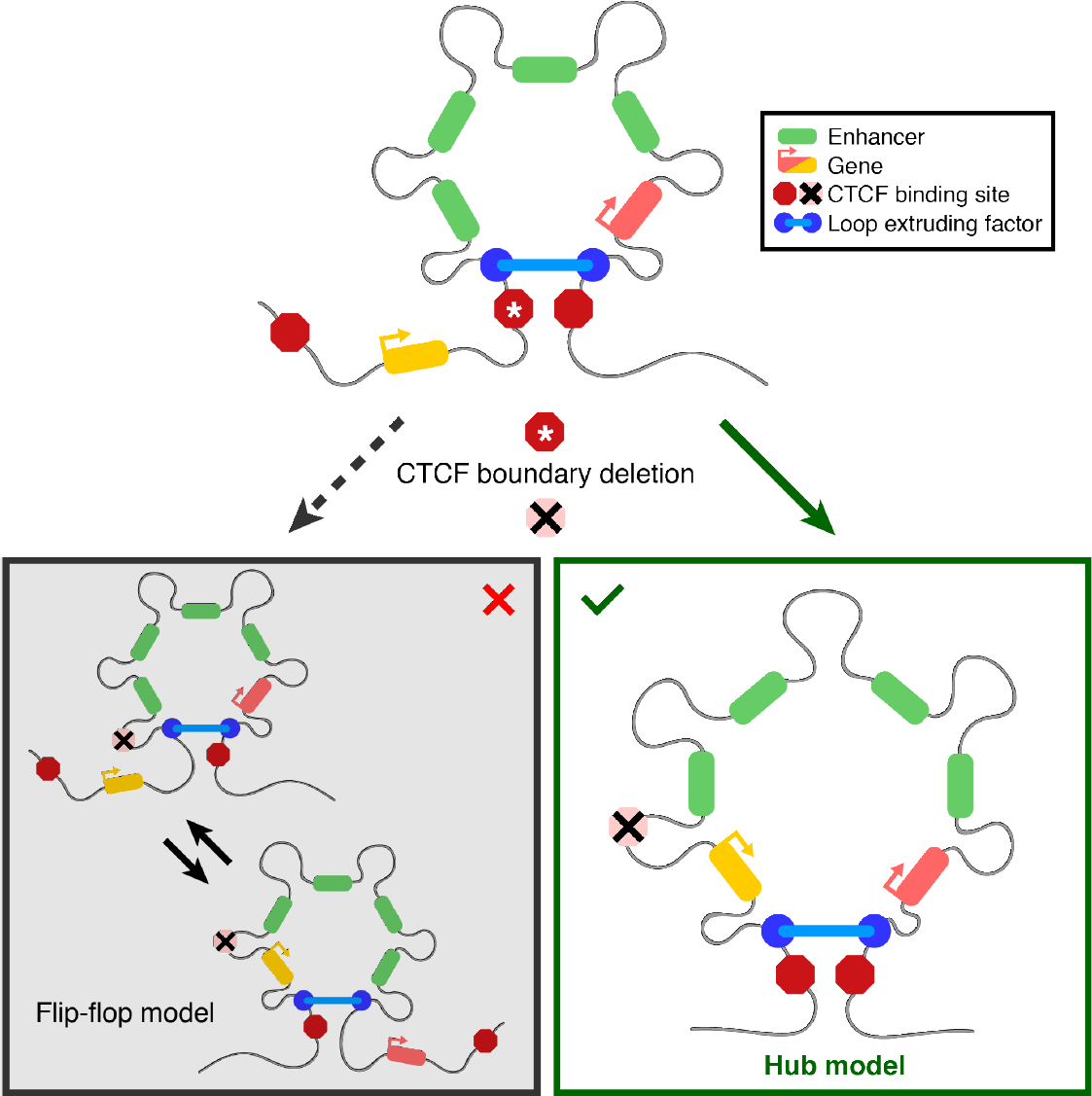
Model of the structural interplay between the regulatory elements in the α-globin locus upon removal of the upstream CTCF boundary. The CTCF boundary upstream of the α-globin enhancers (green) normally constrains their activity to the downstream α-globin genes (red). Removal of this boundary causes upregulation of the genes upstream of the α-globin enhancers (yellow). Our data show that upregulation of these genes is not caused by dynamically switching interactions between the α-globin enhancers and individual promoters (left), but by the formation of a regulatory hub in which all regulatory elements interact simultaneously (right).

Our findings clarify how the activity of strong enhancers is distributed between the multiple genes surrounding these elements. In agreement with previous findings (de Wit et al., 2015; Guo et al., 2015; Hanssen et al., 2017; Lupiáñez et al., 2015; Narendra et al., 2015), enhancers and promoters do not interact beyond strong CTCF-binding sites at TAD boundaries, since removal of the HS-38/-39 boundary is required for the upstream genes to be activated by the α-globin super-enhancer.

By contrast, within a single TAD, all promoters interact with the super-enhancer in a common nuclear compartment. This is consistent with the recent model of transcriptional activation in phase-separated nuclear condensates. However, even in the context of these cooperative structures, the activity of enhancers may not always be distributed equally between all promoters in a TAD. This might partially be explained by the relative position of promoters with respect to the enhancers. Reported examples of promoter competition have often described situations where an active promoter located in between an enhancer and another, more distal promoter causes reduced activity of the distal promoter (Bartman et al., 2016; Cho et al., 2018a; De Gobbi et al., 2006; Wijgerde et al., 1995). It is possible that the proximal highly transcribed gene forms a barrier to loop extrusion (Brandão et al., 2019), due to accumulation of large amounts of transcriptional machinery and regulatory factors, which reduces interactions between the more distal promoter and the enhancers. However, the resulting functional competition does not occur due to mutually exclusive interactions, but because of reduced access to a cooperative condensate in which multiple regulatory elements can interact together.

Interestingly, we have shown that inclusion of the upstream genes in the α-globin hub causes upregulation of their expression, but not to the exceptionally high levels of the downstream α-globin genes (Hanssen et al., 2017). This could be explained by the lower frequency of interaction and inclusion of these genes in the chromatin hub, which likely corresponds to a nuclear condensate. However, it is also possible that biochemical processes within such condensates play a role, which might form another layer of regulation and potential competitive effects. With rapid technological advancements in single cell genomics and imaging, it will be interesting to relate chromatin structures to levels of gene expression in single cells in the future. This could provide important insights into the mechanisms that control gene activity and how mutations that disrupt regulatory chromatin structures contribute to human disease.

## Methods

### Animals and cells

We previously generated the D3839 mouse model, using TALEN and CRISPR-Cas9 to create small 19 bp and 26 bp deletions in the core CTCF-binding motifs of HS-38 and HS-39, respectively (Hanssen et al., 2017). We performed all described experiments in primary cells obtained from spleens of female D3839 or WT C57BL/6 mice treated with phenylhydrazine, and selected erythroid cells based on the erythroid marker Ter119 using magnetic-activated cell sorting, as previously described (Davies et al., 2015). Experimental procedures were in accordance with the European Union Directive 2010/63/EU and/or the UK Animals (Scientific Procedures) Act (1986) and protocols were approved through the Oxford University Local Ethical Review process.

### Capture-C – experimental procedure

We performed Capture-C experiments in 3 biological replicates of primary erythroid cells derived from WT and D3839 mice following the Next-Generation Capture-C protocol (Davies et al., 2015). We used the DpnII restriction enzyme for digestion during 3C library preparation. We designed capture oligonucleotides targeting the DpnII fragments containing the R1 and R2 enhancers using CapSequm (Hughes et al., 2014). Figure 1 shows the interaction profiles from the viewpoint of the R2 enhancer. Data from the R1 viewpoint have been published previously (GEO accession code GSE97871) (Hanssen et al., 2017). The Capture-C libraries were sequenced on the Illumina MiSeq platform (V2 chemistry; 150 bp paired-end reads).

### Capture-C – data analysis

We analyzed Capture-C data as previously described (Davies et al., 2015), using scripts available at https://github.com/Hughes-Genome-Group/CCseqBasicS. Because PCR duplicates are removed during data analysis, Capture-C accurately quantifies chromatin interactions (Oudelaar et al., 2017). The Capture-C profiles in Figure 1 represent the mean number of unique interactions per restriction fragment from 3 replicates, normalized for a total of 100,000 interactions on the chromosome analyzed, and scaled to 1,000. The differential profile highlights the interactions in D3839 cells after subtracting the normalized number of unique interactions in WT cells from those in D3839 cells. Interactions within a proximity zone of 1 kb around the viewpoint and with restriction fragments that were targeted by other capture oligonucleotides in the multiplexed capture procedure were excluded from analysis to prevent artefacts.

### Tri-C – experimental procedure

We performed Tri-C experiments in 3 biological replicates of primary erythroid cells derived from WT and D3839 mice following the protocol available on Protocol Exchange (Oudelaar et al., 2018b). We used the NlaIII restriction enzyme for digestion during 3C library preparation. We added Illumina TruSeq adaptors using NEBNext DNA Library Prep reagents and Ampure XP beads (Beckman Coulter: A63881). We prepared WT libraries using the NEBNext DNA Library Prep Master Mix Set for Illumina (New England Biolabs: E6040S/L) according to the manufacturer’s protocol. For each biological replicate, we performed 2-3 parallel reactions using 6 ug 3C library for sonication and all recovered material (~4.5 ug) for the subsequent library preparation. We amplified each library preparation reaction with a different index, using 2 separate PCR reactions per reaction (a total of 4–6 PCR reactions per biological replicate) to maximize library complexity. This procedure resulted in a total of 7 technical replicates with unique indices. We prepared D3839 libraries using the NEBNext Ultra II DNA Library Prep Kit for Illumina (New England Biolabs: E7645S/L) according to the manufacturer’s protocol. For each biological replicate, we sonicated 4 ug 3C library, after which we split all recovered material (~3 ug) over two parallel library preparation reactions. We amplified the two parallel reactions with the same index using 2 separate PCR reactions (a total of 4 PCR reactions per biological replicate) to maximize library complexity for each biological replicate. This resulted in a total of 3 replicates with unique indices. Because the Ultra II reagents are more efficient than the standard DNA Library Prep reagents, this resulted in comparable complexity for each biological replicate and similar data depth for both conditions (Supplementary Figs. 1 and 2). We pooled all libraries to enrich for our viewpoints of interest using a multiplexed double capture procedure with custom designed capture oligonucleotides (Supplementary Tables 1 and 2). The Tri-C libraries were sequenced on the Illumina NextSeq platform (V2 chemistry; 150 bp paired-end reads).

### Tri-C – data analysis

We analyzed Tri-C data using scripts available at https://github.com/Hughes-Genome-Group/CCseqBasicS and https://github.com/oudelaar/TriC. Briefly, we used the CCseqBasic pipeline (flags: --CCversion CS5 --nla --sonicationSize 700 --wobblyEndBinWidth 6) to perform the initial fastq processing and aligning of the data, filter out spurious ligation events and PCR duplicates, and exclude interactions with restriction fragments that were targeted by other capture oligonucleotides in the multiplexed capture procedure. We used a custom script to select reads with 2 or more reporters to calculate multi-way interaction counts between reporter fragments for each viewpoint. We visualized these interactions in contact matrices at 1 kb resolution, after normalizing for the total counts in each matrix and correcting for the number of restriction fragments present in each bin. We integrated this workflow in the CCseqBasic pipeline, which is available at https://github.com/Hughes-Genome-Group/CCseqBasicS. To allow for direct comparisons between WT and D3839 cells, we scaled all contact matrices to 100 normalized interactions per bin. We derived differential matrices to highlight the interactions specific for each condition after subtracting the normalized interactions in WT cells from those in D3839 cells or vice versa. We also generated regular pair-wise interaction profiles based on the total interaction counts. These ‘Tri-Capture-C’ profiles were derived as described above (Capture-C – data analysis). To calculate the enrichment of multi-way interactions between cis-regulatory elements of interest (highlighted in Figs. 2 and 3), we calculated the counts in the bins in a 2 kb radius surrounding the foci of interest in the matrix and expressed these counts as a percentage of the total number of counts in the matrix. To analyze the differences between the WT and D3839 replicates, we used unpaired, two-tailed t-tests. Statistical analyses were performed with Student’s two-tailed t-tests using GraphPad Prism software.

## Acknowledgements

We thank Matthew Gosden for technical advice and help with next-generation sequencing. This work was supported by Wellcome (Genomic Medicine and Statistics PhD Programme, references 105281/Z/14/Z and 109110/Z/15/Z; Chromosome and Developmental Biology PhD Programme, reference 099684/Z/12/Z; Wellcome Trust Strategic Award, reference 106130/Z/14/Z) and the Medical Research Council (MRC Core Funding and Project Grant, reference MR/N00969X/1). A.M.O. is funded by a Stevenson Junior Research Fellowship at University College, Oxford.

---

**Supplementary Figure 1:**
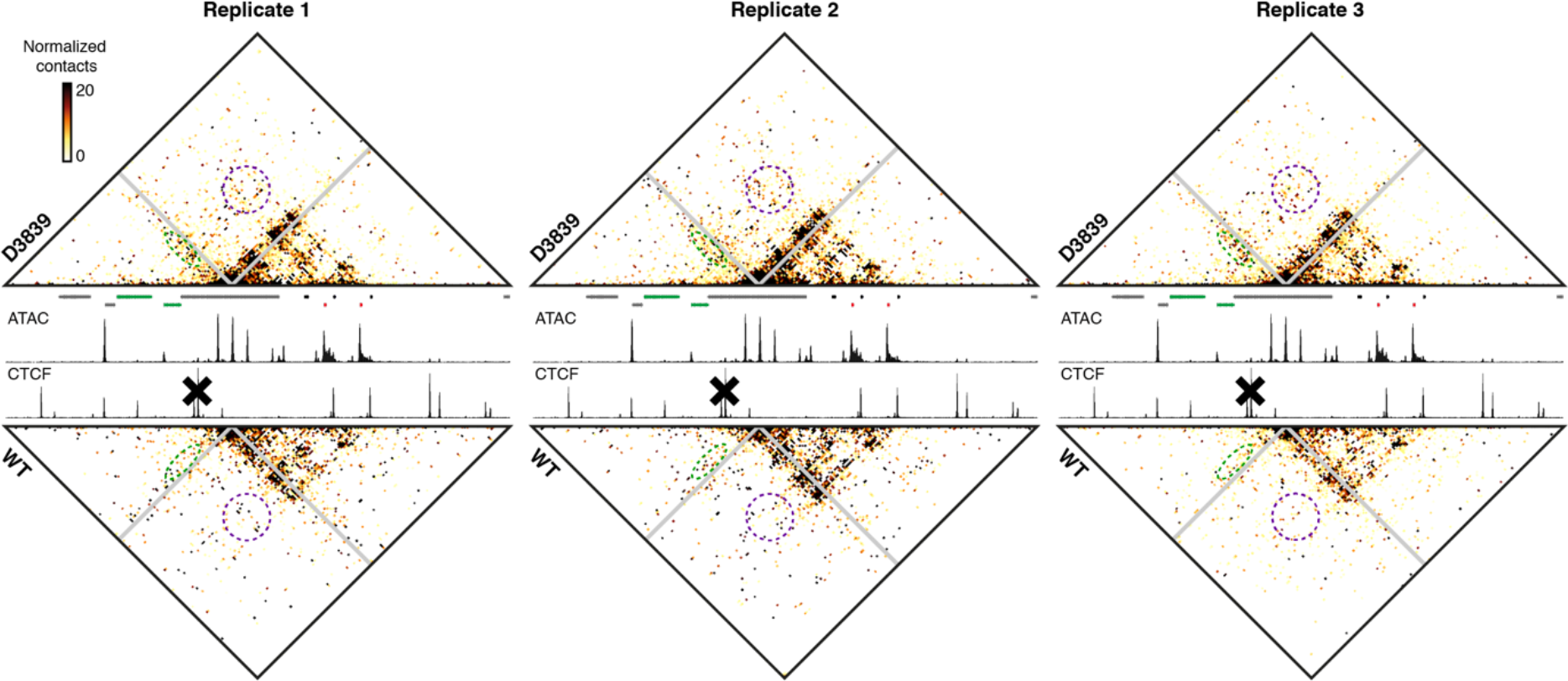
Reproducibility of R2 Tri-C contact matrices. Tri-C contact matrices showing multi-way chromatin interactions with R2 in individual biological replicates of D3839 (top) and WT (bottom) erythroid cells. Matrices represent normalized, unique contact counts at 1 kb resolution with proximity contacts around the R2 viewpoint excluded (gray diagonal). Individual replicates show similar patterns of increased R2 interactions with R1 and the Mpg/Rhbdf1 promoters (green) and the Mpg/Rhbdf1 promoters and the α-globin promoters (purple) in the D3839 cells compared to WT cells. Gene annotation, open chromatin (ATAC) and CTCF occupancy in WT erythroid cells are shown in the middle. Coordinates (mm9): chr11:32,070,000–32,250,000.

**Supplementary Figure 2:**
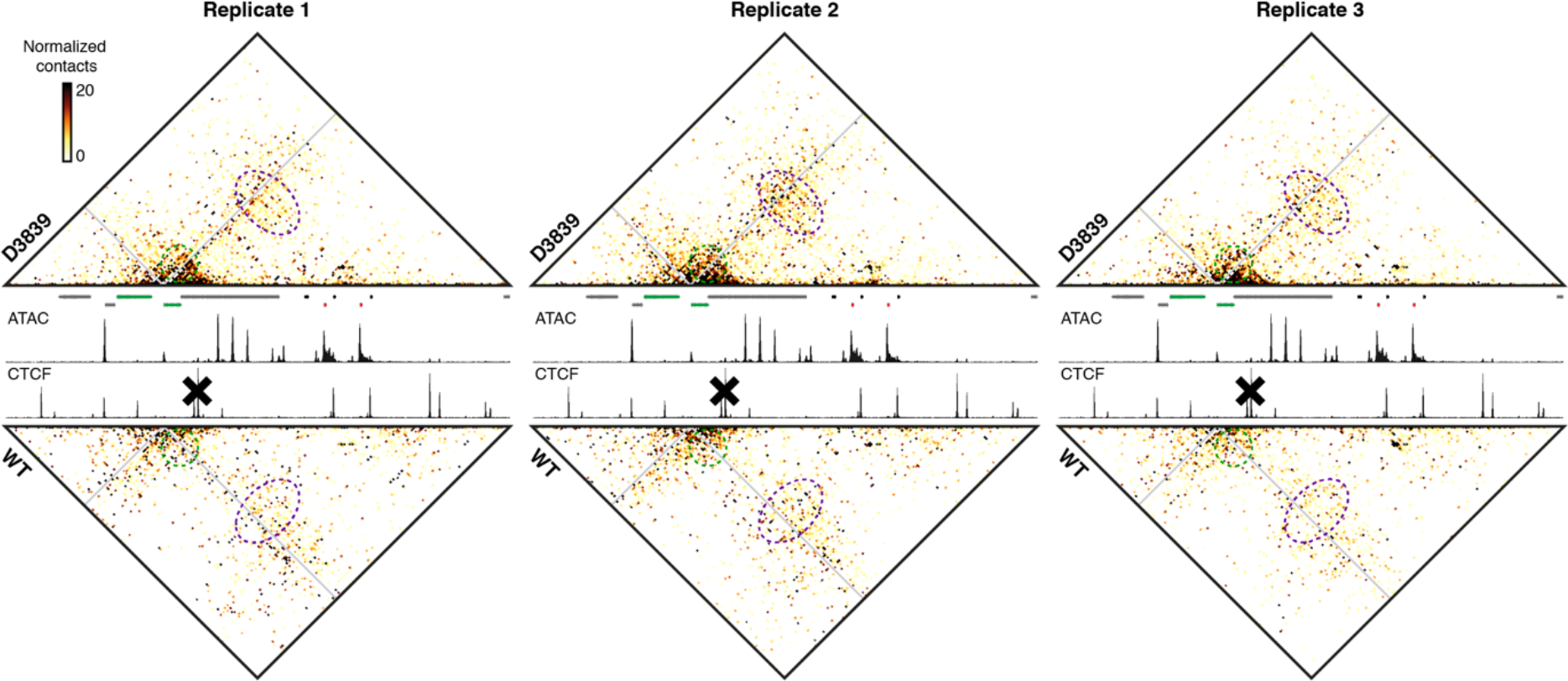
Reproducibility of Mpg Tri-C contact matrices. Tri-C contact matrices showing multi-way chromatin interactions with Mpg in individual biological replicates of D3839 (top) and WT (bottom) erythroid cells. Matrices represent normalized, unique contact counts at 1 kb resolution with proximity contacts around the Mpg viewpoint excluded (gray diagonal). Individual replicates show similar patterns of increased proximal Mpg interactions, including R1 and the Mpg/Rhbdf1 promoters (green) and the Rhbdf1 promoter and the α-globin enhancers/promoters (purple) in the D3839 cells compared to WT cells. Gene annotation, open chromatin (ATAC) and CTCF occupancy in WT erythroid cells are shown in the middle. Coordinates (mm9): chr11:32,070,000–32,250,000.

**Supplementary Figure 3:**
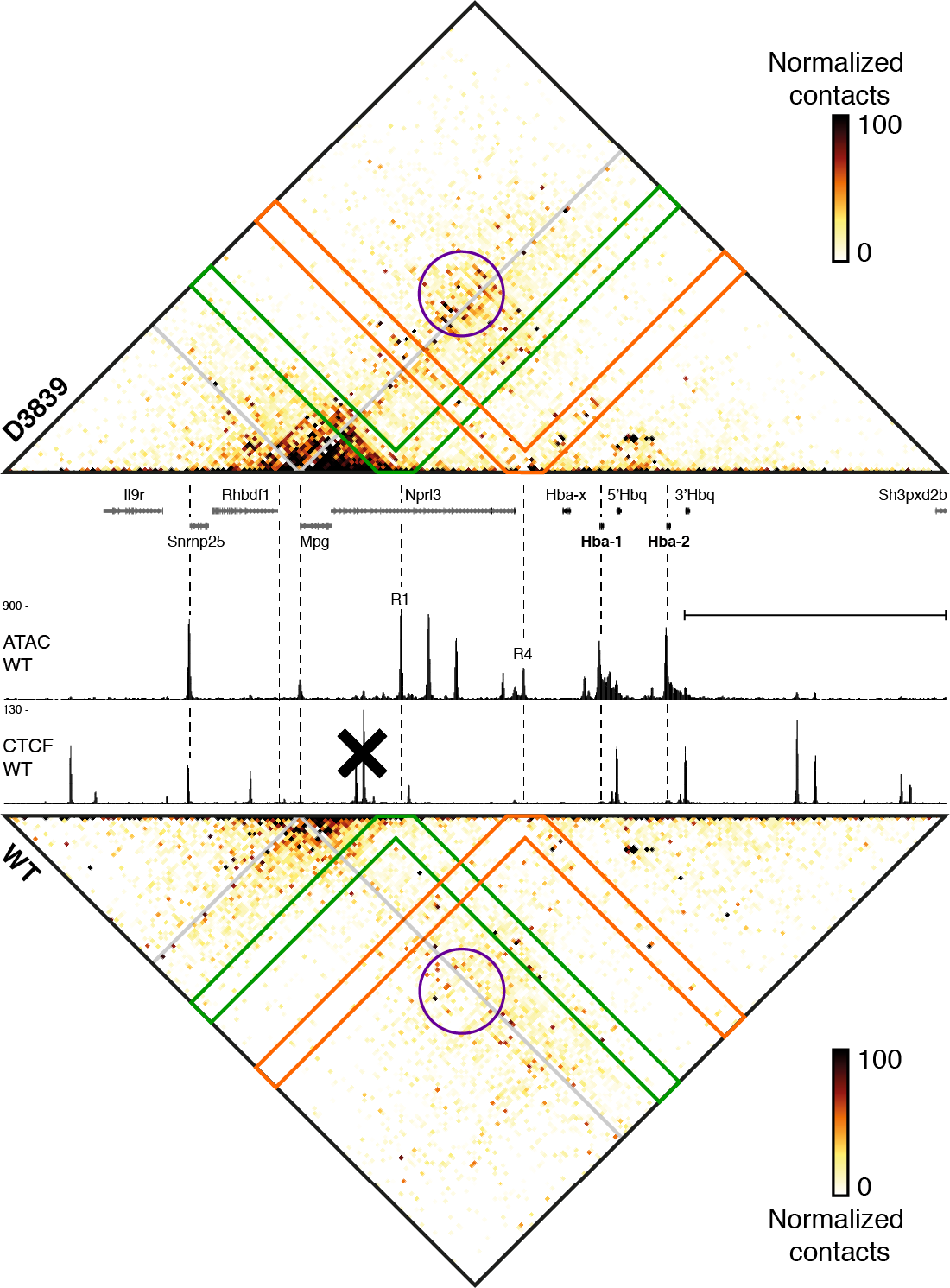
Multi-way interactions with the Mpg promoter. Tri-C contact matrices showing multi-way chromatin interactions with Mpg in D3839 (top) and WT (bottom) erythroid cells. Matrices represent mean numbers of normalized, unique contact counts at 1 kb resolution in 3 replicates with proximity contacts around the Mpg viewpoint excluded (gray diagonal). Gene annotation, open chromatin (ATAC) and CTCF occupancy in WT erythroid cells are shown in the middle. Coordinates (mm9): chr11:32,070,000-32,250,000. To emphasize that the Mpg promoter preferentially interacts with the α-globin enhancers in a complex which includes the α-globin and Rhbdf1 promoters, we have highlighted the regions of the contact matrices that show all the multi-way interactions between Mpg and R1 (green) and between Mpg and R4 (orange). When Mpg interacts with R1 or R4 in D3839 cells, there are clear enrichments over the other α-globin enhancers and the α-globin and Rhbdf1 promoters, indicating that Mpg preferentially interacts with these elements in a complex. The formation of a structure in which multiple promoters interact together is also evident from the increased Mpg interactions with the α-globin and Rhbdf1 promoters (purple) in D3839 cells.

**Supplementary Table 1.**
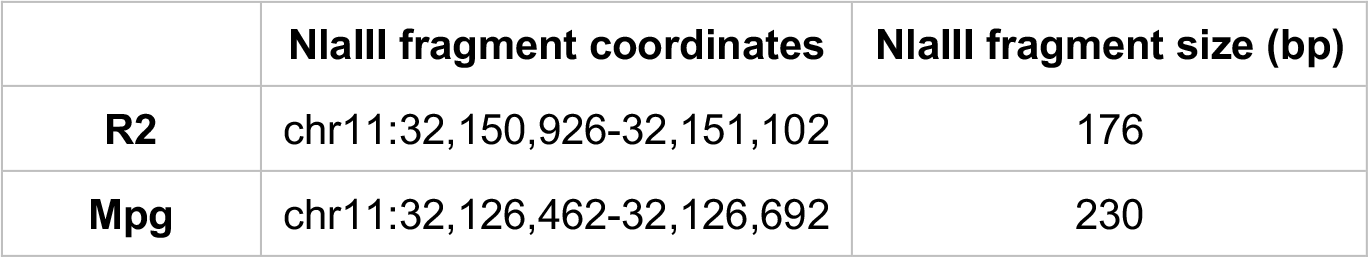
Tri-C viewpoints. Overview of the coordinates and sizes of the restriction fragments used as viewpoints in the Tri-C experiments. The oligonucleotide pools we used for viewpoint enrichment also contained oligonucleotides targeting the following NlaIII fragments: chr11:32137188-32137324, chr11:32100062-32100217 and chr11:32160146-32160318. These restriction fragments were excluded from analysis to prevent artefacts.

|  | **NlaIII fragment coordinates** | **NlaIII fragment size (bp)** |
| --- | --- | --- |
| **R2** | chr11:32,150,926-32,151,102 | 176 |
| **Mpg** | chr11:32,126,462-32,126,692 | 230 |

**Supplementary Table 2.**
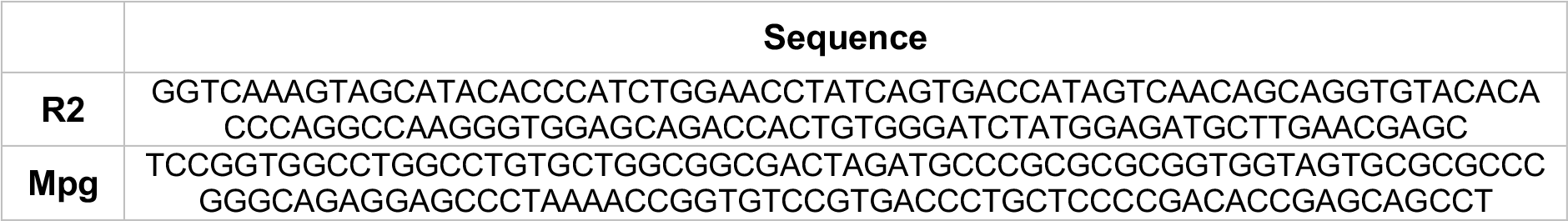
Tri-C capture oligonucleotides. Overview of the sequences of the Tri-C capture oligonucleotides we used to enrich for viewpoints of interest. These 120 bp sequences were designed to target the middle of the restriction fragments listed in Supplementary Table 1.

